# The prefusion structure of the HERV-K (HML-2) Env spike complex

**DOI:** 10.1101/2025.03.19.644160

**Authors:** Ron Shaked, Michael Katz, Hadas Cohen-Dvashi, Ron Diskin

## Abstract

The human endogenous retrovirus K (HERV-K) is a retrovirus that got assimilated into the human genome in ancient times and has been inherited in our germline ever since. It enters cells using a class-I spike protein (Env) that mediates receptor recognition and membrane fusion. On top of having a biological role during development, HERV-K is activated in amyotrophic lateral sclerosis, various cancers, and other pathological conditions. Antibodies that target the HERV-K spike complex have therapeutic value, flagging the spike as a novel drug target. Here, we use cryo-EM to determine the trimeric structure of the HERV-K spike. The spike presents a distinct structure, which substantially differs from other class-I fusogens. Nevertheless, some general architectural features suggest a common origin with other retroviruses. Our structural analysis points to the putative receptor binding sites of the spike and provides insights into its function. The ability to structurally characterize the HERV-K spike may facilitate the development of antibody-based therapies.

**Significance:** Retroviruses integrate their genomes into host cell DNA. When this occurs in germline cells, the retroviral elements can be inherited and become part of the offspring genome. HERV-K is one such retrovirus that integrated into the human genome in ancient times. Over time, it gained a role in embryonic development but is also linked to various diseases, making it a potential drug target. To enter cells, HERV-K uses a spike protein, whose atomic structure we determined using electron microscopy. This structure reveals insights into the spike’s function and HERV-K’s evolutionary ties with contemporary viruses. This structural information further provides a foundation for future drug development.

## Introduction

The human genome contains numerous ancient retroviral segments that integrated in the far history into the germline of our ancestors and, hence, were inherited throughout evolution. When the first draft of the human genome became available, it was realized that up to 8% of our genome is composed of retroviral elements (1). While most of the retroviral genome insertions are shared between humans and apes, dating their insertion to time periods that precede the emergence of humans, one family of human endogenous retroviruses (HERV) known as HERV-K has a subtype called human endogenous MMTV-like (HML) - 2 that is unique to humans (2), making it the most recent one to enter our germline. Due to the sequence similarity with the mouse mammary tumor virus (MMTV), the HERV-K (HML-2) is considered to belong to the betaretrovirus genus of the *Orthoretrovirinae* subfamily within the *Retroviridae* family of viruses (3, 4). Whereas most of the ancient retroviral elements in our genome became inactive due to the accumulation of mutations and recombination events, some retained functional proteins, and there is at least one insertion of HERV-K (HML-2) (out of hundreds of insertions in our genome) that retained full open reading frames for the viral *gag, pro, pol*, and *env* genes (5).

Retaining functional ORFs of these retroviral elements throughout evolution suggests a possible biological role. Indeed, it was discovered that HERV-K (HML-2) plays an important function during embryogenesis (6), but it is typically silenced in healthy adults. Nevertheless, viral elements are activated and produced in a variety of pathologies like cancers (7-12), neuronal motor diseases like amyotrophic lateral sclerosis (ALS) (13, 14), HIV-1infection (15), in addition to senescence (16). In some cases, the specific production of *env*, which is the viral gene encoding the spike (Env) protein of HERV-K, is linked to the pathology and hence makes it a potential target for therapeutic interventions; Endocytosis of the HERV-K (HML-2) spike following attachment to the cellular receptor CD98HC (17), leads to neurotoxicity, which promotes ALS (18, 19). Blocking this entry route with monoclonal antibodies (MAbs) makes a promising new avenue for treating ALS (19, 20). Also, antibodies that target the HERV-K (HML-2) Env, which is abundant in lung adenocarcinoma, were shown to have immunotherapeutic value (21). Hence, the spike complex of HERV-K (HML-2) emerges as a prime target for a variety of human diseases.

The HERV-K (HML-2) spike complex is predicted to be a class-I fusogen that mediates viral entry (3, 22). It is translated as a membrane-embedded precursor protein that trimerizes and requires proteolytic cleavage by furin for activation. Furin cleavage separates the surface subunit (SU) from the transmembrane subunit (TM) (3, 23). The SU domain serves as the receptor binding domain (RBD) and mediates the recognition of the cellular receptors, which in the case of HERV-K (HML-2) were shown to be CD98HC (amino acid heterodimer transporter composed of SLC3A2 and SLC7A5) (17), as well as heparan sulfate (24). The TM domain, which crosses the membrane, makes the fusogenic domain of the spike (3, 23). As a promising new drug target, there is a significant interest in obtaining structural information for the HERV-K (HML-2) spike, but this protein complex has eluded structural determination efforts so far. Here we report the high-resolution EM structure of the HERV-K (HML-2) spike protein. This structure reveals how the HERV-K (HML-2) Env assembles into a functional trimer, points to its putative receptor binding sites, and provides clues for how this trimer may be triggered by acidic conditions. This structural information further informs us about the organization of Env spike complexes within the betaretrovirus genus and reveals some similarities to Env proteins in other genera of the *Retroviridae* family. This structure of the HERV-K (HML-2) spike may also help to promote the development of novel therapeutics that target this spike.

## Results and Discussion

### The overall structure of the HERV-K spike

To determine the structure of the HERV-K (HML-2) spike (referred to as ‘HERV-K’ hereafter), we expressed in HEK293F cells the full-length Phoenix consensus-strain of HERV-K (25) fused to a Flag tag at its C’ end, following a similar methodology that we successfully used for investigating spikes from Arenaviruses (26, 27). We solubilized the spike in detergent, purified it using anti-Flag affinity beads, and determined a three-fold symmetric structure at 2.1 Å resolution using single-particle cryo-EM (Fig. S1, Table S1). The high-quality EM density map allowed us to model the majority of the ectodomain (*i*.*e*., the extracellular portion) of the spike. Disordered regions include a few residues at the termini of SU and the transmembrane/cytoplasmic portions of TM (Fig. 1a). The spike rises more than 114 Å above the surface of the membrane and has a width of about 75 Å, without considering its N-linked glycans (Fig. 1b). The SU domains are separated from one another (Fig. 1b), and a clear solvation layer is formed at the interface between adjacent SU domains (Fig. S2). Near the three-fold symmetry axis of the spike, this separation between the SU domains makes a continuous large cavity that extends down to the TM domain of the spike (Fig. 1b). The SU spans the entire length of the ectodomain; from the apex region where it is the sole constituent of the spike, all the way to the membrane-proximal region (MPER), where its C’ forms a small helical segment that pairs with the TM domain (Fig. 1c). The C’ of SU at the MPER, interacts with the helical stem of TM that is visible up to 10-residues before the beginning of the transmembrane helix (Fig. 1a). Although not modeled, low-resolution EM density that likely extends all the way to the membrane is visible in C1-reconstructed maps of various subclasses (Fig. S3), revealing that the MPER region is helical and is made through the association of three TM-derived helices with the three C’ copies of SU (Fig. 1c). The N’ of SU is solvent exposed, has conformational flexibility in its first 10-residues that were not resolved in the EM density map, and is located near the MPER region of the spike, close to the C’ of SU (Fig. 1c). The N’ of TM, which is visible from the first residue, Phe467, is separated more than 50 Å from the C’ of SU (Fig. 1c). This region of TM, which is mostly hydrophobic, contains the fusion peptide of the spike and is buried at the interface between SU and TM (Fig. 1c, Fig. S4). The side chain of the N-terminal Phe467 interacts with a hydrophobic pocket that is formed on the surface of a three-helical bundle (further discussed below) made by the three symmetry-related TM subunits (Fig. S4). The positively charged amino terminus of TM is accommodated by forming a hydrogen bond with a water molecule that is part of an extensive solvation layer between SU and TM (Fig. S4).

**Figure 1.**
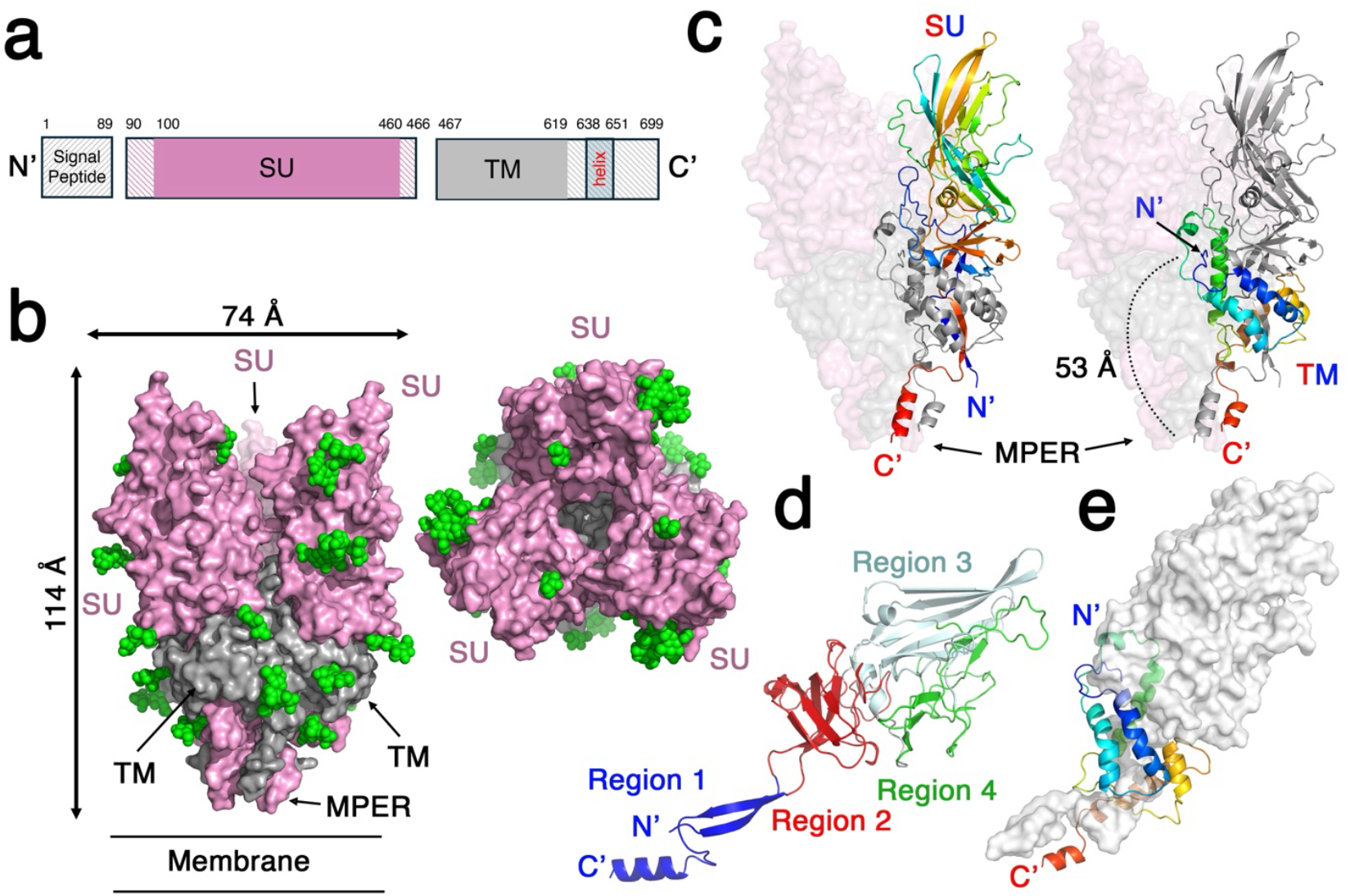
The overall structure of the HERV-K spike complex. **a**. Schematic diagram showing the organization of the HERV-K’s *Env* gene. Areas with solid-colored backgrounds were visible in the EM density map and were modeled. **b**. The trimeric structure of the spike complex. The SU (pink) and TM (grey) domains are shown using surface representation from ‘side’ and ‘top’ views on the left and right sides, respectively. N-linked glycans are shown as green spheres. The relative location of the membrane and the dimensions of the spike are annotated. **c**. The overall organization of the SU and TM domains. The SU (left image) and the TM (right image) domains of a single SU/TM pair are shown as a ribbon diagram, rainbow-colored from the N’ (blue) to the C’ (red), which are annotated. The rest of the spike is shown as a semi-transparent surface for reference. **d**. The specific organization and local folding regions of the SU domain. A single SU domain is shown as a ribbon diagram, colored and annotated according to the four topological regions. **e**. The specific organization of the TM domain with respect to SU. A close-up view of a single TM domain rainbow-colored from the N’ (blue) to the C’ (red). The TM wraps around an SU domain that is illustrated as a semi-transparent white surface.

The SU domain adopts a β-sheet-rich fold that could be divided into four topological regions (Fig. 1d, Fig. S5, and Fig. S6). ‘Region-1’ makes an elongated stem that connects the main body of SU with the MPER region of the spike. It is made of two anti-parallel β-strands, which are donated by the termini regions of SU (Fig. 1d). ‘Region-2’ includes two sandwiched β-sheets that are interconnected with extended loops (Fig. 1d, Fig. S5, and Fig. S6). The most distinctive feature of ‘Region-3’ is a single elongated β-sheet that is partially exposed on one side and is capped on the other side with loops and a couple of small β-strands that make ‘Region-4’ (Fig. 1d, Fig. S5, and Fig. S6). ‘Region-4’ has a small β-sheet that interacts with ‘Region-2’ and a complex structure of loops that wrap around ‘Region-3’ (Fig. 1d, Fig. S5, and Fig. S6).

Searching the PDB using the DALI server (28) identifies only a single structure (Z score of 7.9) that resembles the SU of HERV-K. This structure is of Syncytin-2 (PDB: 7oix) (29), which is part of the ancient human endogenous retrovirus FDR (ERV-FDR) class-I fusogen that utilizes major-facilitator superfamily transporter (MFSD2A) as a receptor and is important in forming a blood-barrier in the placenta by inducing syncytia (29-31). Aligning these two models using TM-align (32) provides a TM-score of 0.41 (normalized to the length of SU from HERV-K), indicating only marginal similarity, as could be appraised visually (Fig. S7). Syncytin-2 mostly aligns with ‘Region-2’ of HERV-K’s SU, but it has an elongated β-sheet that resembles ‘Region-3’ of SU, despite projecting to a different direction (Fig. 1d, Fig. S7). Trying to align a more remote SU domain like the one of the gammaretrovirus Friend murine leukemia virus (33), provides an even lower TM-score of 0.21, indicating very little structural similarity of the HERV-K SU to other retroviruses. The TM domain has an almost exclusive α-helical fold (Fig. 1e, Fig. S8, and Fig. S9). The TM completely wraps around ‘Region 1’ of SU, positioning α-helices and loops all around it (Fig. 1e). Since TM has a simple α-helical fold, the DALI server (28) identifies many remotely similar structures (highest Z score of 4.5 with an E3 ligase), but none of them belongs to an envelope protein.

The relatively high-resolution EM map provided rich structural details, some of which are not always available for class-I fusogens. One important aspect is the pattern of the disulfide bonds of HERV-K, which are all apparent in the map. There is a total of five disulfides that form in the SU domain (Fig. S5, Fig. S10) and one additional disulfide that forms in the TM subunit (Fig. S8, Fig. S10). Interestingly, the SU subunit also includes an unpaired cysteine residue (Cys142) that is in the vicinity of SU’s disulfide number 4 (Fig. S10). Disulfide number 5 in the SU domain of HERV-K has a CXXC pattern (Fig. S5, Fig. S10), which is a reminiscence of the CXXC motif in gamma-type retroviral fusogens (34, 35). However, HERV-K lacks a CX_6_CC motif in the TM domain, nor does it form a linking disulfide between the SU and TM domains, which is a hallmark of gamma-type retroviral fusogens (36). Besides disulfides, other interesting aspects of the spike that are visible in the EM map include an extensive solvent layer (1626 modeled water molecules), which provides information about the solvation and accommodation of charged/polar species at the core of the protein (Fig. S2). Interestingly, on top of a total of three pre-proline peptide bonds at a *cis* configuration that each SU domain has, there is also a rare non-pre-proline *cis* peptide bond forming between Gly317 and Glu318 of the SU domain that facilitates a sharp turn of the main chain (Fig. S11). Another unusual structural feature is a fully buried salt bridge forming between Arg269 and Asp265 in the SU domain (Fig. S12). These two residues make an important stabilizing factor in this region of SU since Asp265 further binds to two main-chain amine groups, and Arg269 also engages with two main-chain carbonyls (Fig. S12).

### The trimeric organization of the spike

The trimeric organization of the HERV-K spike complex has some distinct features from many other known viral spike complexes. One noticeable feature is the lack of any significant interactions between the three SU domains. While many indirect water-mediated polar interactions form between the SU domains (Fig. S2), only two noticeable direct contacts can be observed; The first one is a polar interaction between Lys224 and a main-chain carbonyl on a neighboring SU, and the second is a Van der Waals interaction forming by clustering of all three symmetry-related Leu453 residues at the spike’s three-fold symmetry axis (Fig. 2a). Hence, the trimeric configuration of the HERV-K spike can be almost entirely attributed to TM-TM interactions (with only a minor contribution of SU-TM contacts in the context of the trimer, see below), which are indeed substantial; Analyzing the pair-wise TM-TM interactions reveal that between each pair of chains there is a total of 2933 Å^2^ buried surface area (1506 Å^2^ on the first chain and 1427 Å^2^ on the other), which sums to 8799 Å^2^ buried surface area for the formation of the trimer.

**Figure 2.**
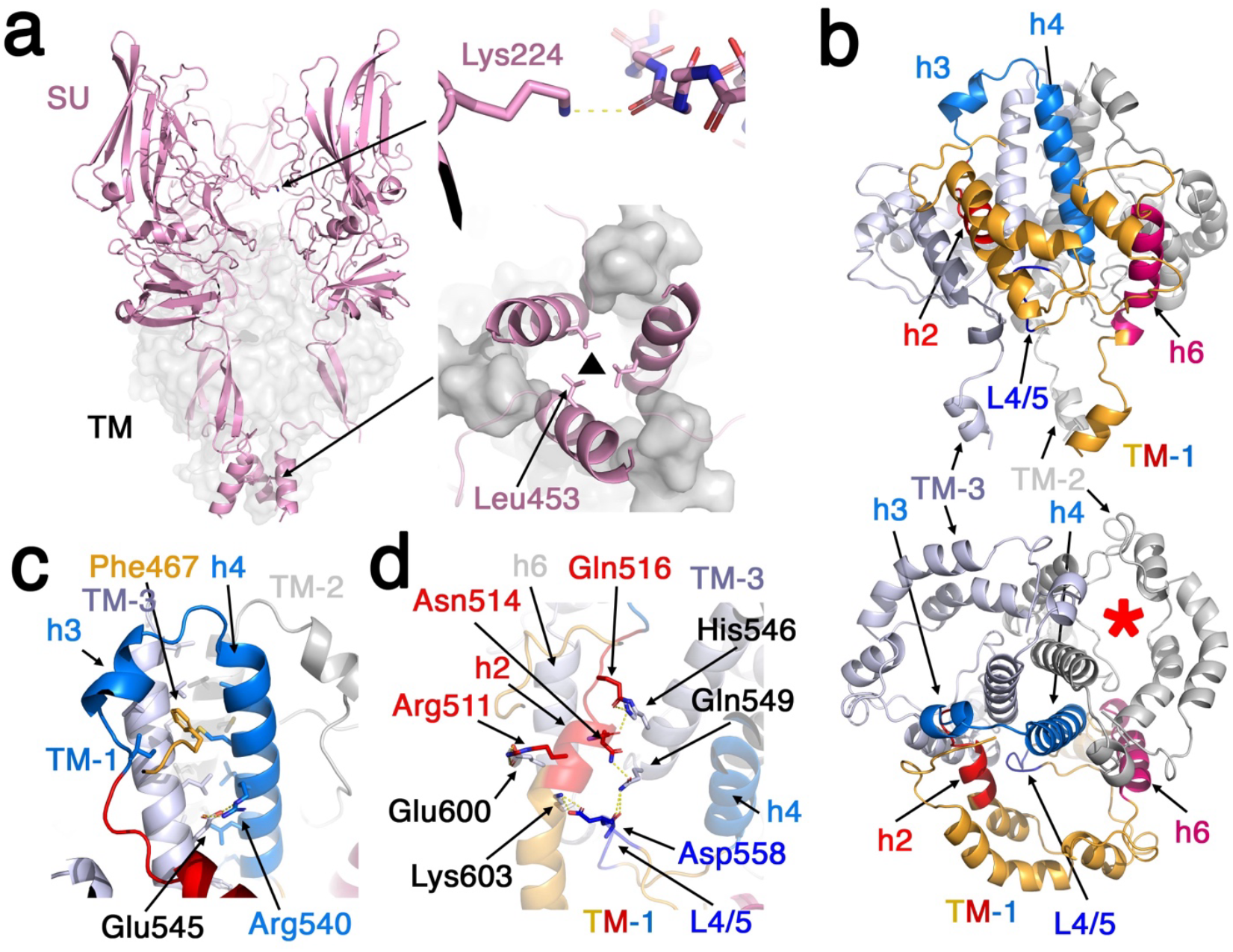
The trimeric organization of the HERV-K spike complex. **a**. SU-SU interactions are scarce. The SU domains are shown as pink ribbons, and the TM domains are shown using a semi-transparent grey surface representation in an illustration of the entire spike from a ‘side’ view. The direct contact points between the SU domains are pointed, and zoom-in views are shown on the right. The lower zoom-in image is from a ‘bottom’ view of the spike. **b**. Overview of the TM-TM interactions. The three TM domains are shown as ribbons and colored in orange, grey, and livid for TM-1, TM-2, and TM-3, respectively. The three TM domains are shown from a ‘side’ view (upper image) and from a ‘top’ view (lower image). The interacting regions in TM-1 are differentially colored and annotated. A red asterisk marks the area inside TM that accommodates SU. **c**. A close-up view of the central coiled-coil and the role of h3 and h4 helices in its formation. **d**. The various polar interactions between the h2 helix and loop L4/5 on one TM and the h6 and h4 helices on a neighboring TM.

The interfaces between the TM domains span two distinct regions and involve multiple parts of TM (Fig. 2b). At the heart of the spike, a large coiled-coil is formed by three copies of h4 helices (Fig. 2b, Fig. S8). This coiled-coil configuration is driven by multiple hydrophobic residues along the h4 helices that point toward each other (Fig. 2c). In addition, some polar interactions like a salt bridge between Arg540 and Glu545 from a neighboring h4 helix, further stabilize this coiled-coil (Fig. 2c). The short h3 helix, which just precedes h4, is interacting with the outer side of the central coiled-coil, further expanding the TM-TM interface (Fig. 2b, Fig. 2c). The interaction of h3 with h4 helices of the neighboring TM creates the hydrophobic pocket that accommodates the N-terminal Phe467 of TM (Fig. 2c, Fig. S4). The second distinct interface between the TM domains is primarily hydrophilic. It involves the h2 helix and L4/5 loop on one TM that engages with the h6 helix on the neighboring TM (Fig. 2b, Fig. 2d). This interface consists of multiple salt-bridge and hydrogen bond interactions (Fig. 2d).

### The formation of SU-TM pairs in the HERV-K spike

The TM domain wraps around the two antiparallel β-strands that make ‘Region 1’ of the SU domain (Fig. 1d, Fig. 1e), creating a very substantial interface between them. The buried surface area at this interface reaches a total of 6625 Å^2^ (3361 Å^2^ on SU and 3264 on Å^2^ TM). To accommodate SU, the TM folds such that it creates a cavity (Fig. 2b). This cavity has the central coiled-coil on one side (Fig. 2b) and has a clamp-like formation that grips SU in place using two ‘jaws’ (Fig. 3a). Each ‘jaw’ is made from topologically proximal regions of TM; The first is made from h1 and h2 helices, and the second is made from h5 and h6 helices as well as the L4/5 and L 5/6 loop regions (Fig. 3a). Hence, SU is not topologically tied within TM, and a relative motion of the ‘jaws’ that will increase the gap between h1 and h5 helices of TM, may facilitate dissociation of SU.

**Figure 3.**
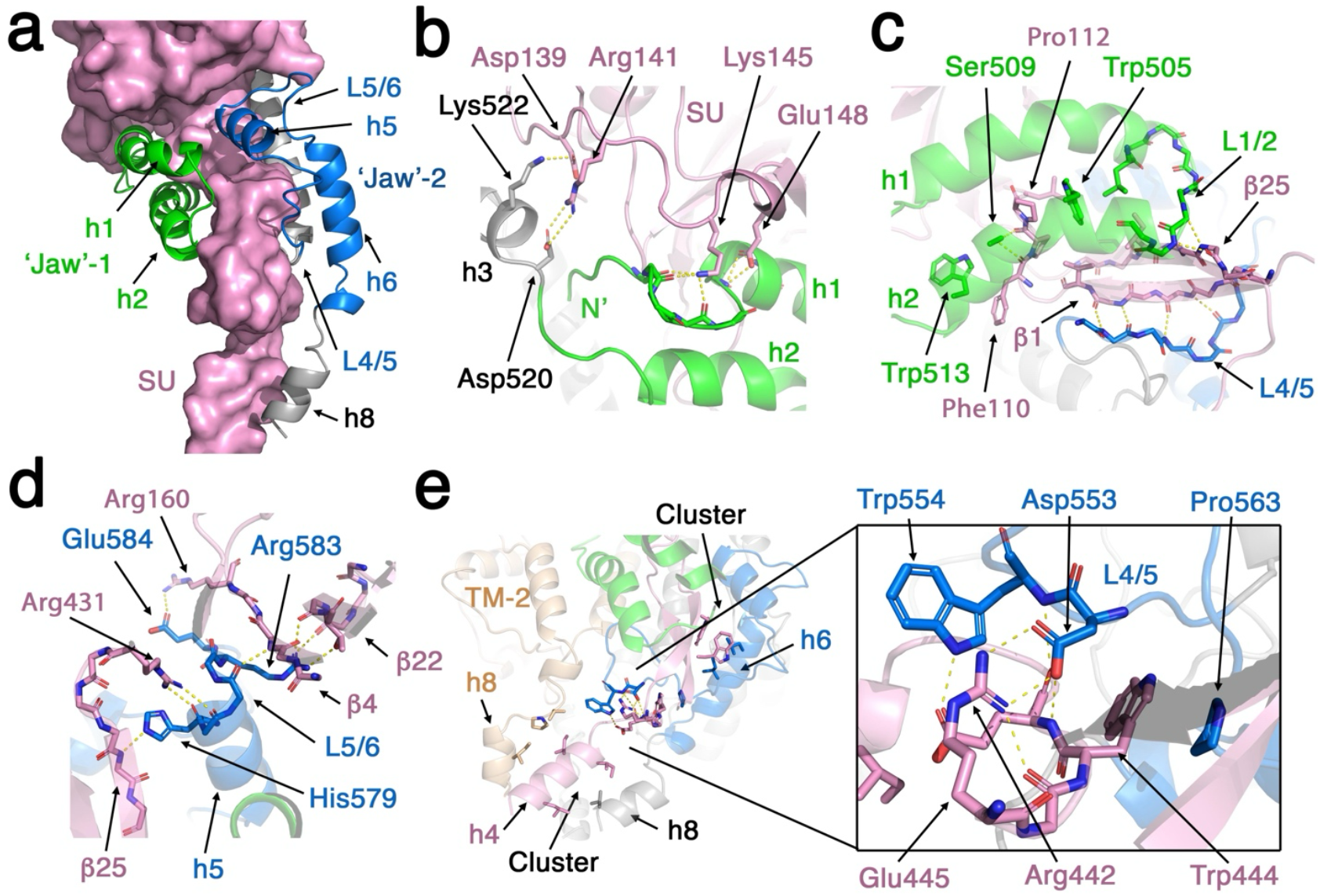
TM binds SU in a clamp-like configuration. **a**. A single TM/SU pair is shown. The SU is represented as a pink surface, and the TM is shown using a ribbon representation and is colored in grey, green, and blue for the main body, the first and the second ‘jaws’ that grip SU, respectively. **b, c & d**. Detailed description of the TM-SU interactions. The same coloring scheme is used as in ‘a’. Polar interactions are illustrated with yellow-dashed lines. **e**. Detailed view of the TM-SU interactions near the MPER region. The image on the right shows a zoom-in view of the indicated region.

The SU-TM interface is highly elaborate and consists of Van der Waals-based hydrophobic interactions, hydrogen bonds, and salt-bridge interactions. Nonetheless, it seems that polar interactions dominate this interface. At the base of ‘jaw’-1, there is a cluster of polar interactions that include two salt bridges between the SU residues Asp139 and Arg141 that pair with Lys522, and Asp520 from the TM domain, respectively (Fig. 3b). In addition, the SU Lys145 satisfies three exposed main-chain carbonyls of TM, which belong to the N-terminal fusion peptide, and Glu148 from the SU domain serves as an N’-cap for h1 helix of TM (Fig. 3b). Further along ‘jaw’-1, there is a cluster of hydrophobic residues that include Trp505 and Trp513, which primarily interact with Phe110 and Pro112 on SU (Fig. 3c). The tip of ‘jaw’-1 that is made of a short L1/2 loop, forms hydrogen bonds with β25 of SU, partially blocking one of the exposed edges of the ‘Region 1’ stem (Fig. 1d, Fig. 3c, Fig. S5). The second ‘jaw’ of TM also makes a myriad of contacts with SU; The L4/5 loop forms a short β-strand that pairs with β1 of SU to satisfy its main chain polar groups with hydrogen bonds (Fig. 3c). The h5 helix of ‘jaw’-2 helps to block the edge of β25 by placing His579 that serves as a hydrogen-bond acceptor for a main chain amine group (Fig. 3d). From the L5/6 loop, Glu584, Arg583, and a main chain carbonyl pair with Arg160, a couple of exposed main-chain carbonyls, and a main-chain amine group of SU, respectively. Interestingly, the SU domain further helps to stabilize TM by donating Arg431, which serves as a C’-cap for the h5 helix of TM (Fig. 3d). The h6 helix of ‘jaw’-2 mostly contributes hydrophobic residues that form a hydrophobic cluster with SU-derived tryptophan residues (Fig. 3e). Toward the MPER region of the spike, L4/5 loop of TM makes multiple contacts with a loop that just precedes the terminal-h4 helix of the SU domain (Fig. 3e). These interactions include another salt bridge between Asp553 of TM with Arg442 of SU, which are also perfectly positioned to form hydrogen bonds with two main chain polar groups of SU (Fig. 3e). Pro563 of TM is forming a stacking-like hydrophobic interaction with Trp444, and Trp554 of TM engages in a hydrogen bond with Glu445 from SU (Fig. 3e). At the MPER region of the spike, which is made of three h4 helices of SU and three h8 helices of TM, a hydrophobic cluster is forming (Fig. 3e). In this region, a neighboring TM makes TM-SU contacts, which also contribute, albeit in a limited manner, to the formation of the trimer (Fig. 3e).

### The putative receptor binding sites of HERV-K

HERV-K can utilize two distinct receptors for cell entry. The first receptor is heparan sulfate (HS) (24), which is a linear glycosaminoglycan made of repeating disaccharide units with various sulfation patterns. A common sulfated disaccharide in HS is made of glucuronic acid (GlcA) and N-*sulfo* glucosamine (GlnNS) (37), which yields a highly acidic polymer. To gain insights into the potential binding site of HS, we analyzed the electrostatic potential of the HERV-K spike (Fig. 4a). The entire apex of the spike has a weak positive surface potential, which may facilitate long-range attraction between HS and HERV-K (Fig. 4a). However, the side of the spike has a continuous patch of a strong positively charged surface potential that also makes a shallow groove (Fig. 4a). Manually fitting a short stretch of a polymer made from GlcA-GlnNS repeating units, indicate that this groove is large enough to accommodate HS (Fig. 4b). Interestingly, unlike the apex of the spike that is partially blocked by N-linked glycans, this side groove of HERV-K is completely exposed (Fig. S13), increasing the likelihood that this patch is involved in binding HS. The positive charge in this patch results from three arginine residues that pose their guanidino groups on the surface of SU (Fig. 4c). Analyzing the conservation of SU using the ConSurf server (38, 39) reveals that Arg168 is a relatively variable residue (Fig. 4d). Compared with Arg168, Arg204 and Arg206 are more conserved, so do other nearby surface residues that contribute for the formation of this putative HS binding groove on the surface of SU (Fig. 4d). Noteworthy, the entire exposed surface of SU is less conserved than its stem, which interacts with TM (Fig. 4d).

**Figure 4.**
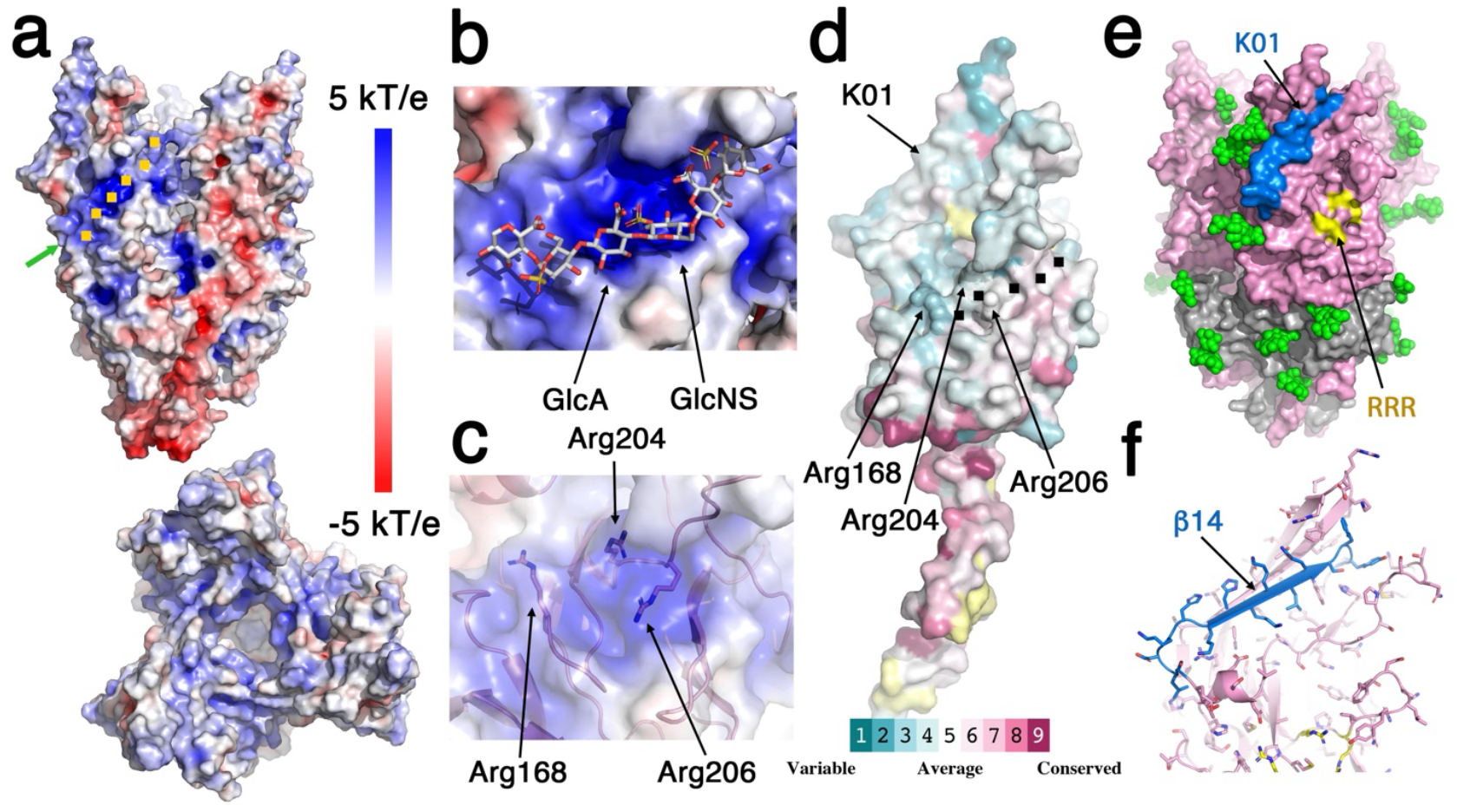
Putative receptor binding site of the HERV-K spike. **a**. Surface electrostatic potential of the spike shown from a ‘side’ view (upper image) and from a ‘top’ view (lower image). The electrostatic potential is illustrated by the color (red to blue gradient represents −5 kT/e to 5kT/e). A basic patch on the surface of SU is noted with a green arrow and traced with orange squares. **b**. Putative accommodation of heparan sulfate inside the basic patch of SU. A short segment made of glucuronic acid (GlcA) and sulfated glucosamine (GlcNS) was manually fitted into the model and is shown using a grey stick illustration on top of the surface that is colored by the electrostatic potential. **c**. Three arginine residues contribute to the basic patch. The residues are shown as sticks underneath a semi-transparent surface and are noted. **d**. Conserved residues in the SU domain. A single SU domain is shown using a surface representation that is colored according to the relative sequence conservation as calculated by the ConSurf server (38, 39) (Red to blue represents conserved to variable). The three arginine residues are noted as well as the basic patch that is traced with black squares. **e**. The HERV-K spike is shown using surface representation. N-linked glycans are shown as green spheres. The linear epitope of MAb K01 is colored blue. The three arginine residues that make the basic patch are noted in yellow for reference. **f**. A close-up view of the linear epitope of MAb K01 on the SU domain. The epitope is colored blue.

The second receptor for HERV-K is CD98HC (17). The interaction with CD98HC leads to neurotoxicity, which could be rescued using an anti-HERV-K MAb called ‘K01’ (19). This antibody recognizes a linear epitope (SLDKHKHKKLQSFYP) that is part of SU. Mapping this epitope on the surface of the trimer reveals that it is located on the side, near the apex of the trimer (Fig. 4e), not very far from the putative binding site for HS. The epitope includes β-strand 14 and some flanking residues (Fig. 4f, Fig. S5), which is located on ‘region-3’ of the SU domain (Fig. 1d). Since MAb K01 interferes with the binding of HERV-K to CD98HC, we can assume that the binding site for this receptor is in the vicinity of K01’s epitope. The epitope and its close vicinity have an average conservation (Fig. 4d), which is similar to that of the putative HS binding site. Noteworthy, the face of SU that contains the K01’s epitope and the putative HS binding site lacks N-linked glycans (Fig. 4e), which is expected for receptor binding sites that need to stay exposed.

### Potential pH-sensors in the HERV-K spike

Following attachment to cells via HS, HERV-K can facilitate membrane fusion in response to acidic pH (Fig. 5a). The obvious and most likely residues that can serve as pH sensors and induce pH-dependent triggering of membrane fusion are histidine residues. Each SU domain has nine histidine residues (Fig. 5b, Fig. S5), and each TM domain has six histidine residues (Fig. 5b, Fig. S8). Many of these histidine residues are either surface-exposed or engage in interactions that are not likely to be affected by protonation. Nevertheless, our structural analysis indicates two histidine residues in the TM domain that their protonation will force a change in their interaction mode. The first one is His579, which is located on the ‘jaw’-2 of TM (Fig. 5b, Fig. 3d). This histidine forms a hydrogen bond with a main chain amine group of SU (Fig. 3d). In this configuration, His579 acts as a hydrogen-bond acceptor, and when protonated, this interaction will no longer persist. Moreover, acquiring a positive charge following protonation may cause electrostatic repulsion from the positively charged Arg431 of the SU domain, which is very close by (Fig. 3d). The second histidine is His546, which is located on the h4 helix that is part of the coiled coil at the center of the spike (Fig. 5c, Fig. 2b). On the one side, His546 interacts with two carbonyls on an adjacent TM domain (Fig. 5c). These are a main-chain carbonyl and a carbonyl from the side-chain of Gln516. For this interaction, His546 is serving as a hydrogen-bond donor. On the other side, His546 engages with Lys595, which is located on a h6 helix from the cognate TM domain (Fig. 5c). For this interaction, His546 is serving as a hydrogen-bond acceptor. When protonated, His546 will no longer be able to engage with Lys595 and will likely repel it due to the electrostatic charge. Interestingly, the h6 helix is part of the TM’s ‘jaw’-2 (Fig. 3a). Hence, the protonation of both His546 and His579 is predicted to weaken the interaction of TM’s ‘jaw’-2 with SU. Importantly, while compelling, testing this hypothesis will be difficult, if not impossible, due to the important structural role of both His546 and His579 in stabilizing the native structure of the HERV-K spike, as is often the case for such pH sensors.

**Figure 5.**
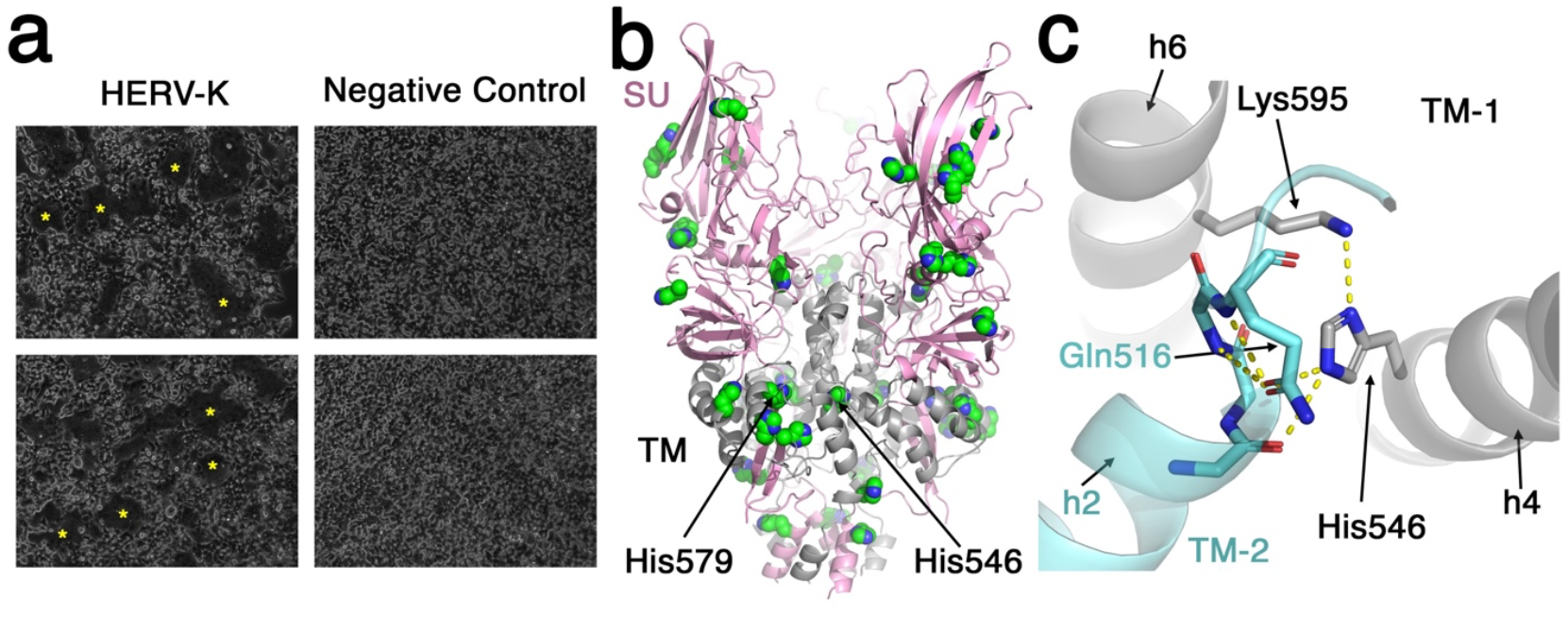
Acidic-induced triggering of membrane fusion. **a**. Syncytia formation assay in cells that express either HERV-K (left) or unrelated protein (negative control, right) following exposure to low-pH buffer. Yellow asterisks mark some of the syncytia. For each condition, two representative images are shown. This experiment was repeated twice. **b**. The histidine residues in the HERV-K spike. All histidine residues are shown as green spheres. His546 and His 579 are indicated. **c**. A closeup view of His546. Two TM domains are shown using grey and cyan ribbon representations. Polar contacts are indicated with yellow dashed lines.

### HERV-K in comparison to other retroviral spike complexes

While structural information is available for some isolated SU domains and for TM domains in a post-fusion state for various retroviral spikes (29, 33, 40, 41), structural information for complete retroviral Env trimers at a pre-fusion state is scarce, with the noticeable exception of HIV-1 from the lentivirus genus, for which there is ample structural information. The only non-lentivirus retroviral spike that its structure is currently available is of a foamy virus (FV) that belongs to the *Spumaretrovirinae* subfamily of the *Retroviridae* (42). Interestingly, the spike complex of FV resembles that of F-type spikes from the *Pneumoridae* family, which is evolutionary very far from the *Retroviridae* (42).

Comparing the HERV-K spike to the Env spike complex of HIV-1 in a pre-fusion closed state (PDB: 5CJX) (43) reveals, at first glance, very different structures (Fig. 6a). The gp120 receptor binding domains of HIV-1 (equivalent to SU in HERV-K) intimately interact with each other at the apex of the spike (Fig. 6a). The gp41 fusion domain of HIV-1 (equivalent to TM of HERV-K) is mostly helical, as TM in HERV-K, but seems to be organized differently (Fig. 6a). However, a closer look at the spikes reveals some organizational features that are preserved in both spikes; Both of the spikes are organized around a central coiled-coil that is formed by three TM/gp41 helices (Fig. 6b). The SU domain and the gp120 domain, although adopting very different structures, have similar topological organization as seen in SU (Fig. 1d), with four distinct regions (Fig. 6c). In both spikes, the fusion subunit engages with the receptor binding domain by engulfing two anti-parallel β-strands that makes ‘region-1’ (Fig. 6c). In both spikes ‘region-2’ of the receptor binding domains is a β-sheet structure that makes extensive contacts with the fusion subunit (Fig. 6c). While pointing to different directions, ‘region-3’ in both spikes is an extended β-sheet structure that is supported by ‘region-4’, which has a more complex organization that includes a β-sheet, loops and helical elements. Hence, the structural information strongly suggests that HIV-1 and HERV-K had a common ancestral origin. This observation further implies that perhaps the entire *Orthoretrovirinae* subfamily shares the same basic architecture for their Env spike complexes. However, to make more definite conclusions, additional structural information for other spike complexes from the various genera of the *Retroviridae* family will need to be obtained.

**Figure 6.**
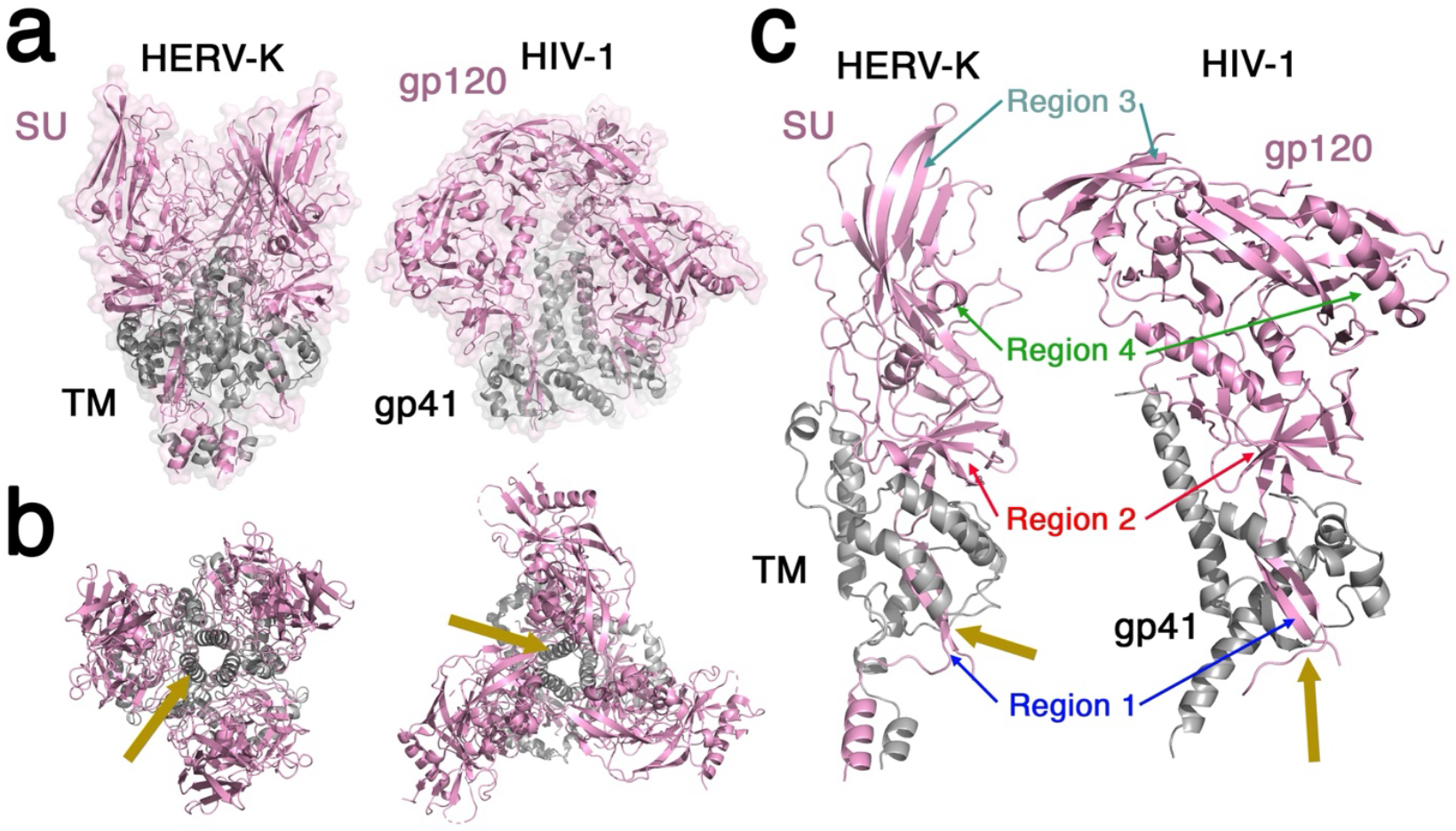
The HERV-K spike shares structural features with the spike of HIV-1. **a**. The HERV-K spike (left) is shown next to the spike of HIV-1 (right, PDB:5CJX), using a ‘side’ view. In both spikes, the receptor binding domains (SU and gp120, for HERV-K and HIV-1, respectively) are colored pink, and the fusion domains (TM and gp41, for HERV-K and HIV-1, respectively) are colored grey. The structures are shown using a ribbon diagram and a semi-transparent surface representation. **b**. The HERV-K and HIV-1 spikes are shown from a ‘top’ view. Orange arrows point to the central coiled coil that is formed by TM/gp41 helices. **c**. SU/TM and gp120/gp41 pairs are shown. The four topological regions of HERV-K (as in Figure 1d) are noted, as well as their equivalent regions in HIV-1.

### Translational implications

The HERV-K Env is emerging as an immunotherapeutic target for various pathologies. Defining exact binding modes and mechanisms of action for MAbs is, in many cases, an important step toward their development for clinical use. The ability to isolate and determine the structure of the HERV-K Env, as we demonstrated here, may facilitate structural studies of the HERV-K spike in complex with MAbs of interest.

## Acknowledgments

The Diskin lab is supported by research grants from the Ernst I. Ascher Foundation, Ben B. and Joyce E. Eisenberg Foundation, Estate of Emile Mimran, Jeanne and Joseph Nissim Center for Life Sciences Research, Donald Rivin, Stanley and Tanya Rossby Endowment Fund, as well as from the Israel Science Foundation (grant No. 209/20). This study was supported in part by a grant from the Moross Integrated Cancer Center.

## Data availability

The EM maps and coordinate file for the HERV-K spike were deposited at the EMDB and PDB, respectively, and are available under accession codes: 9NND/EMD-49572.

## Materials and Methods

### Cloning of Expression Vectors

The full-length HERV-K (HML-2) (Phoenix consensus HERV-K sequence) gene was a gift from Prof. Sean Whelan’s Lab and was subcloned using NotI-KpnI restriction enzymes into pCDNA3.1 with a FLAG tag at the C-terminus.

### expression and purification

HERV-K *env* was expressed in HEK293F suspension cells (Invitrogen). Cells were grown in FreeStyle Medium (Life Technologies), to a density of ∼1.0 × 10^6^ cells per ml before transfection. Cells were transfected using 40 kDa polyethylenamine (PEI-MAX) (Polysciences) at 1 mg ml^−1^, pH 7.0, with DNA at a ratio of 1:2 (DNA:PEI solution). The HERV-K-expressing cells were collected 48 hours post-transfection by centrifugation at 700 × g, 4 °C for 5 min. Membranes were resuspended in a cold lysis buffer (10 mM Tris-HCl pH 8.0, 100 mM MgCl2, 1 mM EDTA, 100 μM phenylmethyl sulfonyl fluoride (PMSF) and 15% v/v glycerol) and homogenized for 5 min on ice. The lysis mixture was centrifuged at 21,000 × g for 25 min, 4 °C. The supernatant was discarded, and pellets were dissolved in a solubilization buffer (20 mM Tris-HCl pH 8.0, 150 NaCl, 10 mM ZnCl_2_, 15% v/v glycerol, 0.1% w/v CHS, 1% w/v LMNG) and homogenized for 10 min on ice. The solubilization mixture was then incubated while rotating for 1.5 hours and centrifuged at 370,000xg for 25 min, 4 °C. The supernatant of this solubilization step was then incubated overnight with 50 μl of Sigma M2 anti-Flag beads (Sigma Aldrich). The anti-Flag beads were then spun down (800 × g, 2 min) and washed by subsequently decreasing amounts of glycerol to 1%, LMNG to 0.14%, and CHS to 0.003%. The protein was eluted with 100 μl of the same buffer of the last washing step supplemented with 0.40 mg/ml 1 × Flag peptide (Genescript) for 1 hour on ice. Eluted protein aliquots were flash-frozen in liquid nitrogen. For western blot analysis, anti-Flag primary antibody (Cell Signaling Technology) was used at a 1:1,000 dilution, followed by horseradish peroxidase (HRP)-conjugated anti-Rabbit secondary antibody (Jackson) at a 1:10,000 dilution.

### Cryo-EM image acquisition and 3D reconstruction

A total of 3.5 μl of purified HERV-K envelope sample was applied on glow-discharged (8 s, 12 mA; Pelco easiGlow, Ted Pella) graphene oxide Quantifoil copper grids, R1.2/1.3, (Electron Microscopy Sciences) using a Vitrobot system (Thermo Fischer/FEI) (3.0 s blotting time, 4 °C, 100% humidity). Samples were incubated on the grid for 1 min before blotting was carried out. Cryo-EM data were collected on the Titan Krios microscope (FEI) operated at 300 kV, using a Gatan K3 direct detection camera. The beam size was 705 nm diameter (fringeless illumination), the exposure rate was 18 e s^−1^ pixel^−1^, and movies were then obtained at 105,000× magnification with a pixel size of 0.824 Å. The nominal defocus range was −0.8 to −2.0 μm. Data processing was carried out with the cryoSPARC v4.1.2 suite (44). Patch motion correction and patch CTF estimation were carried out using cryoSPARC Live. Particles were extracted using a 256-pixel box size, and the data sets were cleaned and classified, as illustrated in Figure S1. Working map were obtained from non-uniform refinement, imposing C3 symmetry, and resolution-based local filtering.

### Model building, refinement, and Structural analysis

A model of HERV-K was generated, based on the EM map, using ModelAngelo (45). This model was then manually completed and refined using Coot (46) and real-space refinement in Phenix (47). Structural analysis and representation were conducted using PyMol (48), ChimeraX (49), and CCP4 (50). Sequence-related secondary structure and topology diagrams were generated using PDBsum (51). Surface electrostatic potential was calculated using APBS tools (52). Conservation analysis was done using the ConSurf server (38, 39). Structural homologs were searched using the DALI server (28), and structural alignment was calculated using TM-align (32).

### Syncytia formation assay

HEK293T cells were seeded onto a 6-well plate (Greiner) at 700K/well and incubated overnight at 37 °C, 8% CO2. After 24 hours, the cells were transfected with lipofectamine 2000 (Invitrogen) at a ratio of 2.5 µl of reagent to 1 µg DNA in Opti-MEM 1x (Gibco). The transfection mixture was then applied to the cells in DMEM (Gibco) and incubated for 4.5 hours, after which point the media was replaced with “Full media”, i.e. DMEM supplemented with 10% Fetal Calf Serum (Gibco), 1% Pen-Strep solution (v/v) (Sartorius), 2.0 mM L-glutamine solution (Biological Industries), and incubated for 48 hours. Afterward, the cells were incubated for 20 minutes in acidified Full media at pH 5.0 (containing 40 mM MES; Sigma Aldrich). The acidified media was then aspirated and replaced with Full media to incubate for an additional 2 hours. Imaging was then carried out on live cells using an Olympus IX51 fluorescence microscope with cellSens Entry software (Olympus LS).

