## Supplemental Information for "The prefusion structure of the HERV-K (HML-2) Env spike complex"

This file contains:

Supplementary Figures: 1-13

Supplementary Tables: 1

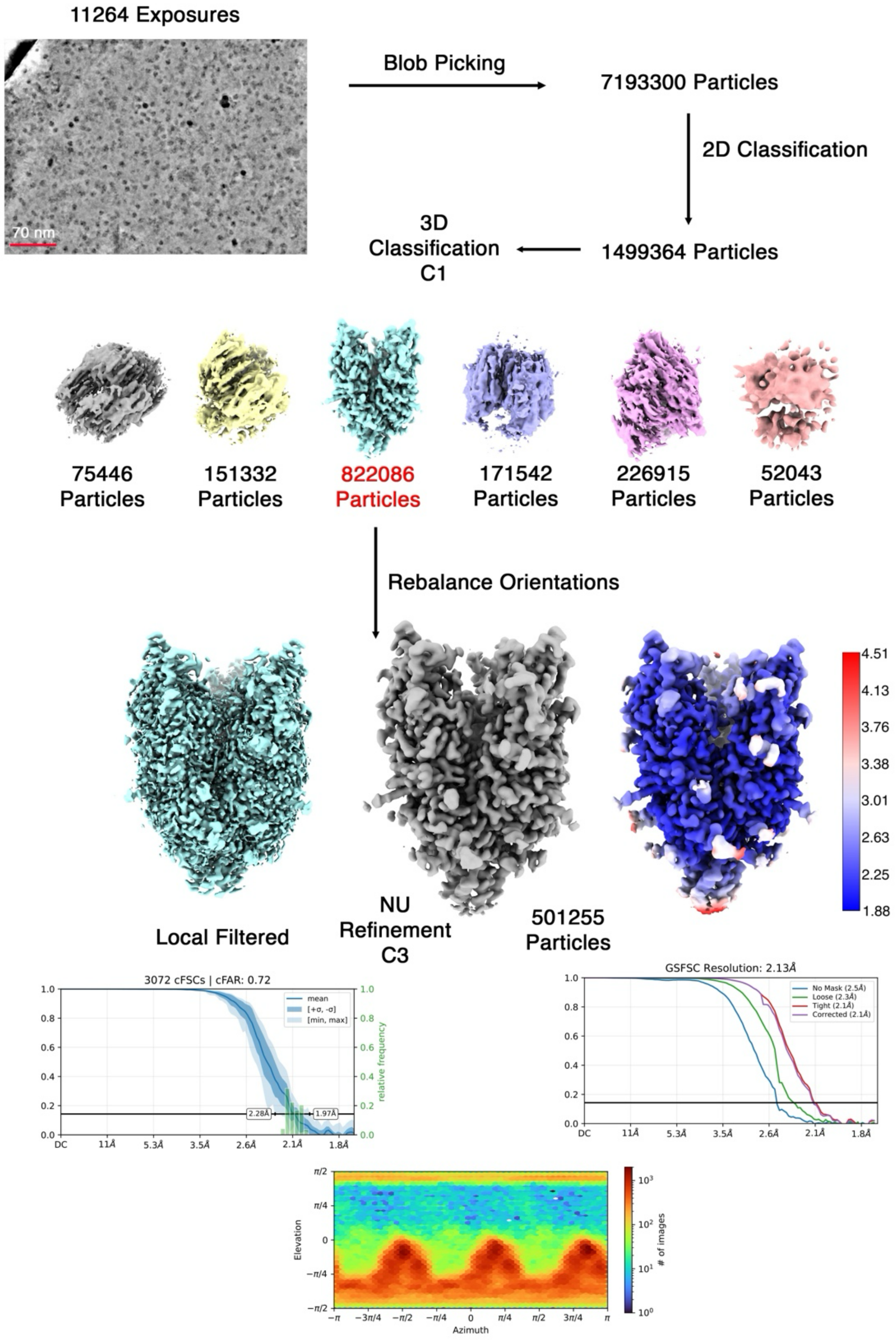


Figure S1 – **The single particle cryo-EM reconstruction pipeline**. The sample micrograph is denoised.

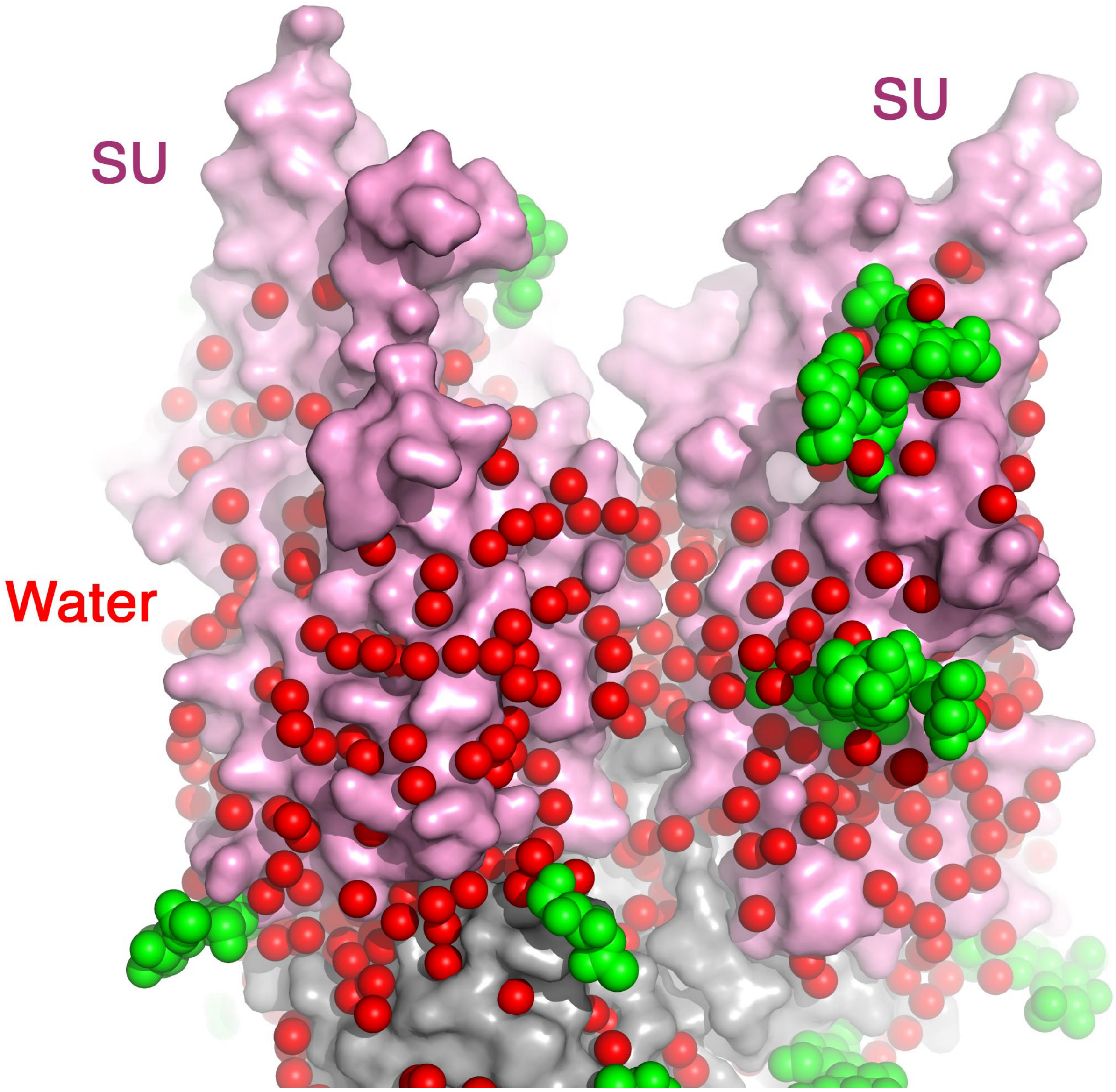


Figure S2 – **The SU domains are separated by a solvation layer**. The front of the spike is shown using a “side view”. Two SU domains are shown with a pink surface representation. Ordered water molecules are shown as red spheres. N-linked glycans are shown using green spheres.

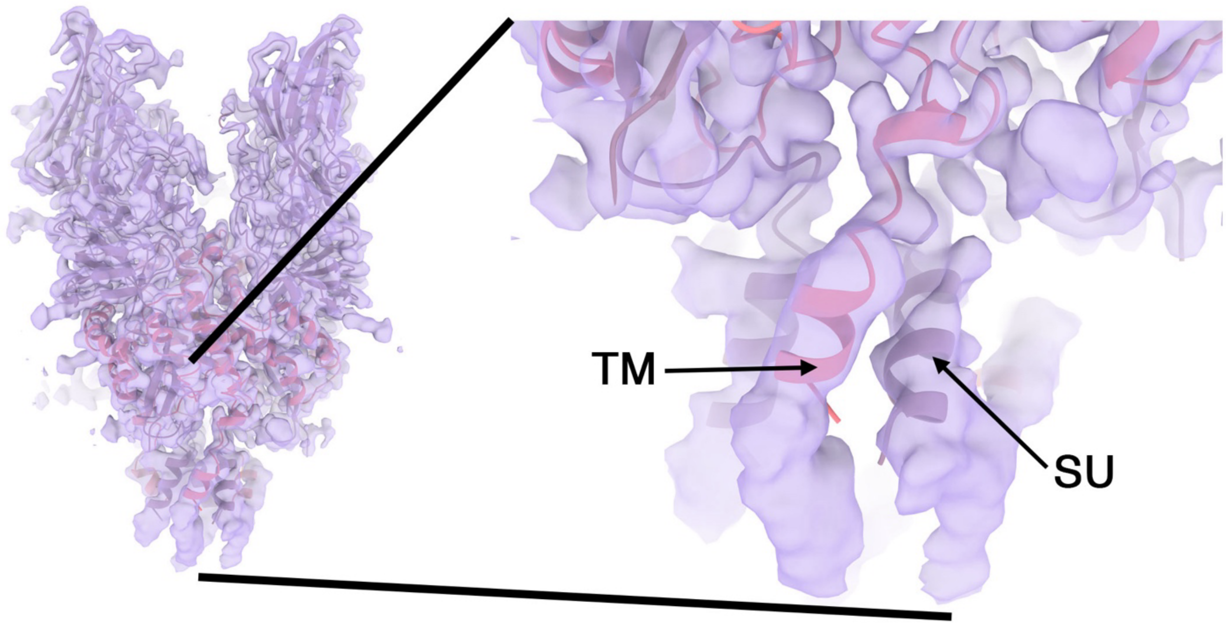


Figure S3 – **The MPER region of the spike is helical**. A C1 reconstructed map of a 3D class using a symmetry-expanded set of particles at map level=0.1 is shown as a semi-transparent surface. The model of the HERV-K spike is shown as a ribbon diagram. The image on the left shows the entire spike, and the image on the right shows a zoom-in view of the MPER region. Additional density beyond the modeled C-termini of SU and TM is visible.

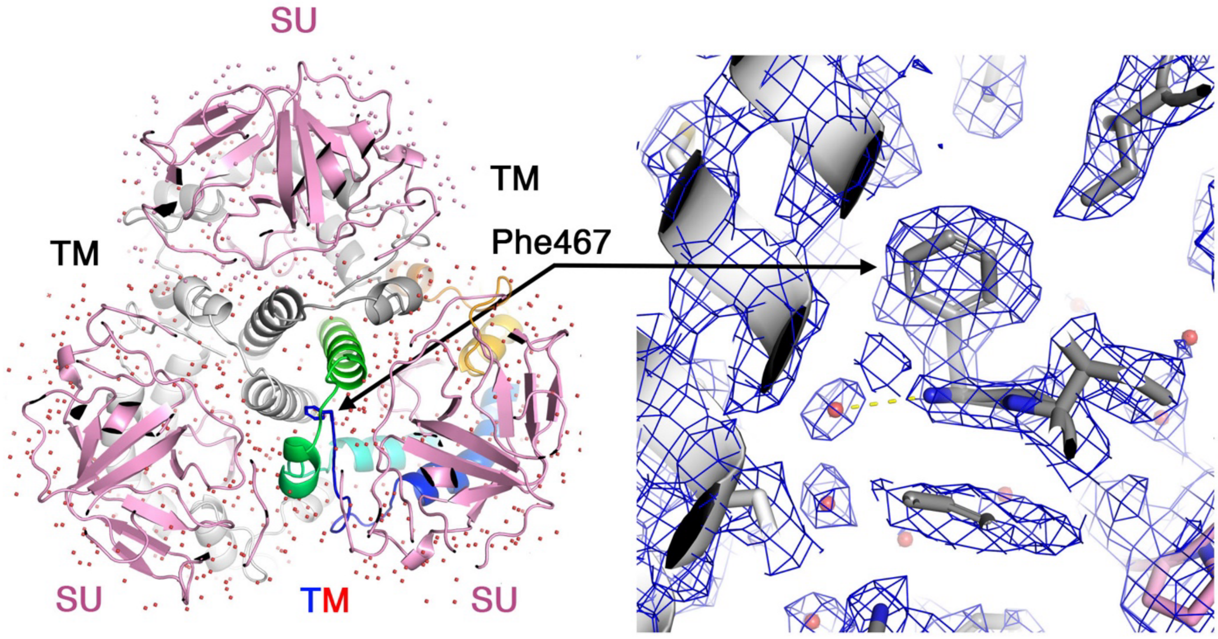


Figure S4 – **The N' of TM is buried at the SU/TM interface and is solvated**. The left image shows the spike from a 'top' view. One TM subunit is rainbow-colored (N' blue, C' red). Water molecules are shown as red spheres. The N-terminal Phe467 interacts with a hydrophobic cavity formed by three TM helices that are organized around the three-fold symmetry axis of the spike. The right image shows a close-up view of Phe467. The EM density is shown as a blue mesh (at σ=5.0). A hydrogen bond between the terminal amine group of Phe467 and a water molecule is illustrated with a yellow dashed line.

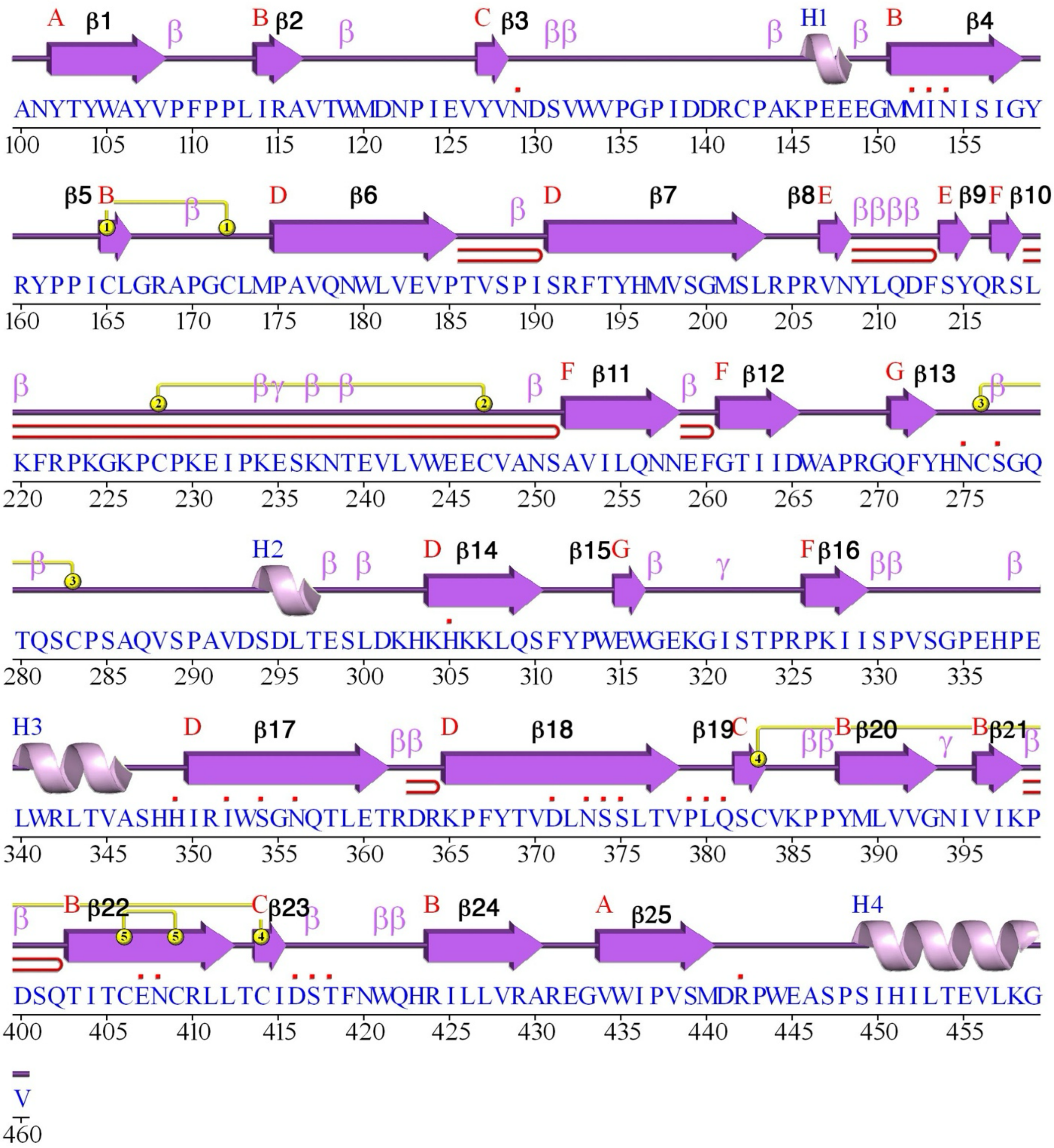


Figure S5 – **Secondary structure of the SU domain**. This diagram was generated using PDBsum (1).

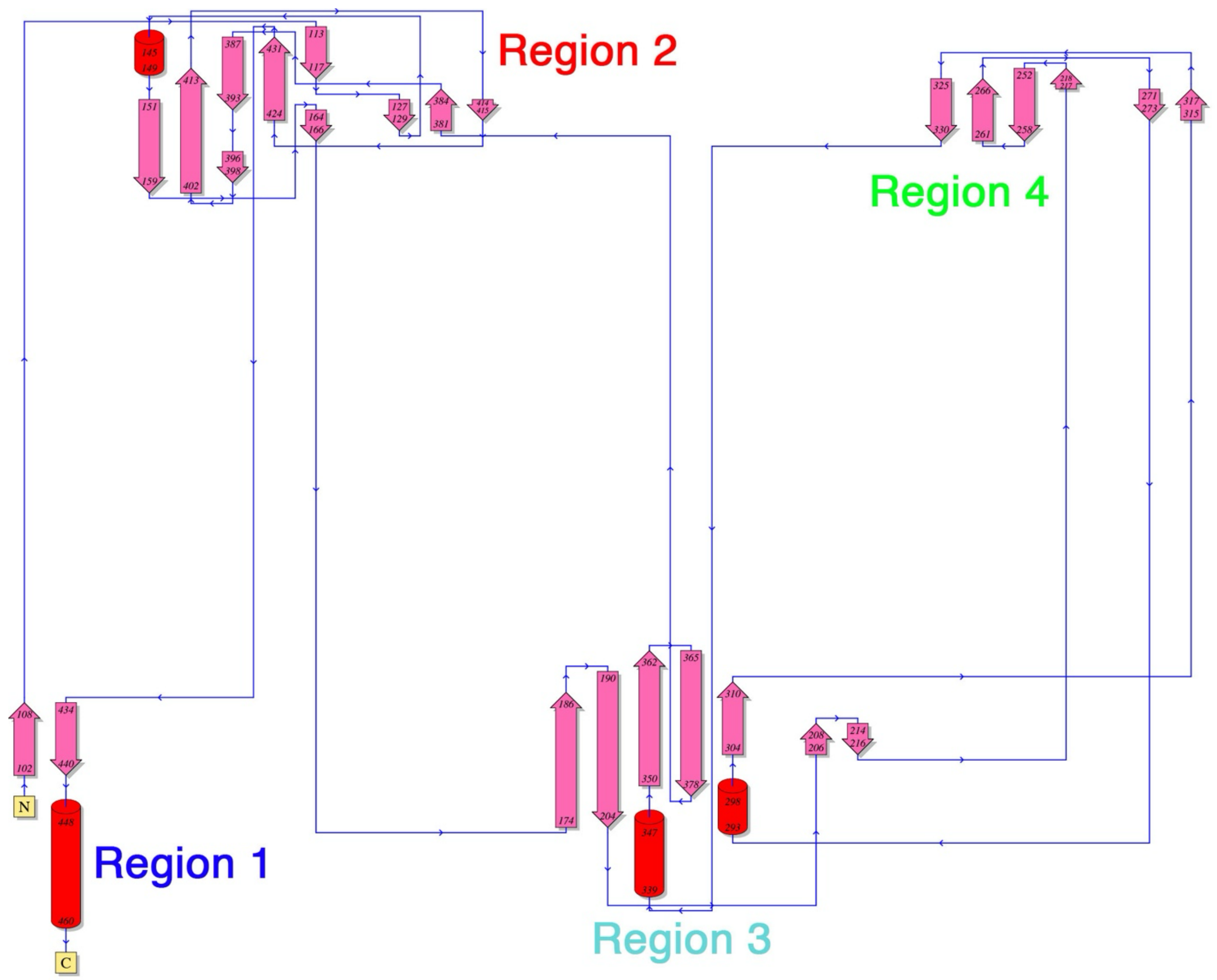


Figure S6 – **The topology of the SU domain**. This diagram was generated using PDBsum (1). The four regions are colored as in Main Figure 1d.

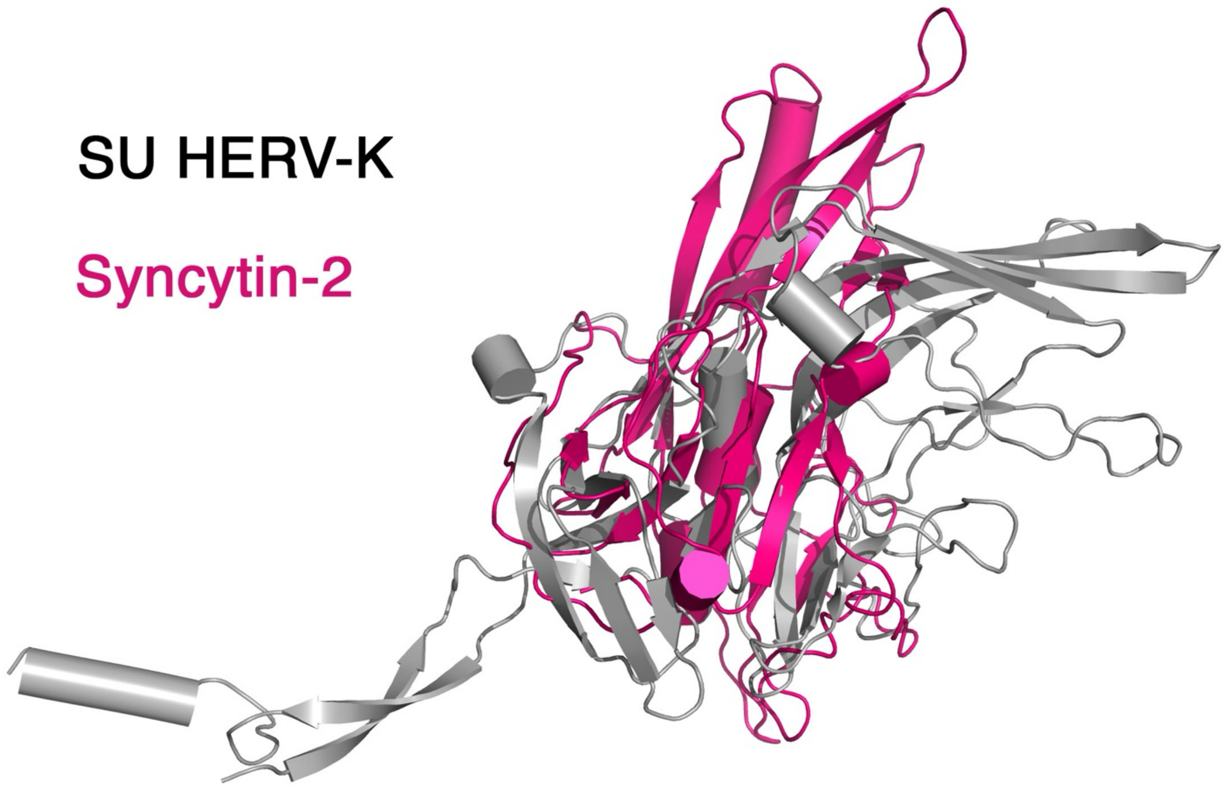


Figure S7 – **Alignment of SU from HERV-K and Syncytin-2 (PDB: 7oix) by TM-align**. The TM-align server calculated RMSD is 4.62 Å based on 195 amino acid pairs that have a distance lower than 5.0 Å.

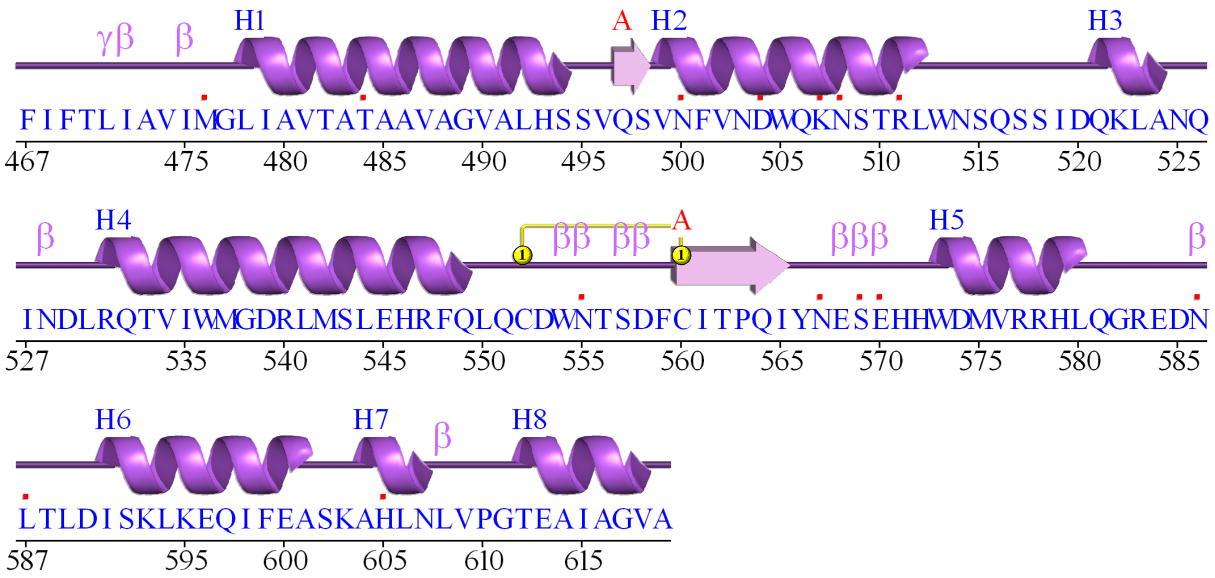


Figure S8 – **Secondary structure of the TM domain**. This diagram was generated using PDBsum (1).

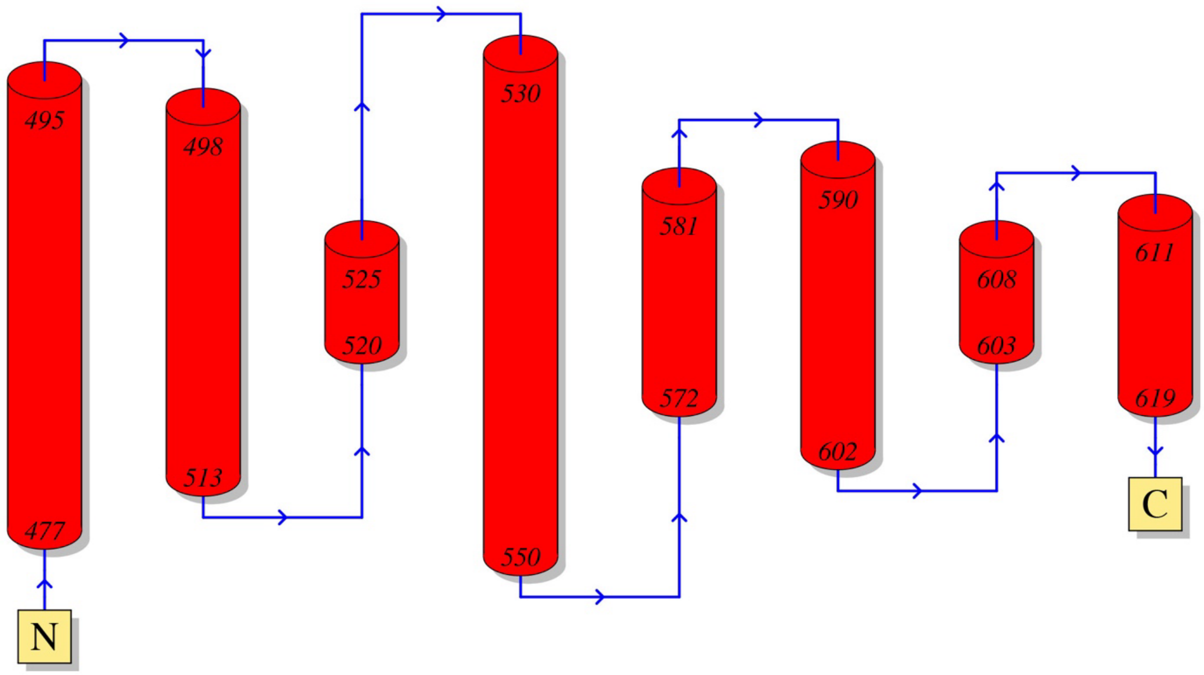


Figure S9 – **The topology of the TM domain**. This diagram was generated using PDBsum (1).

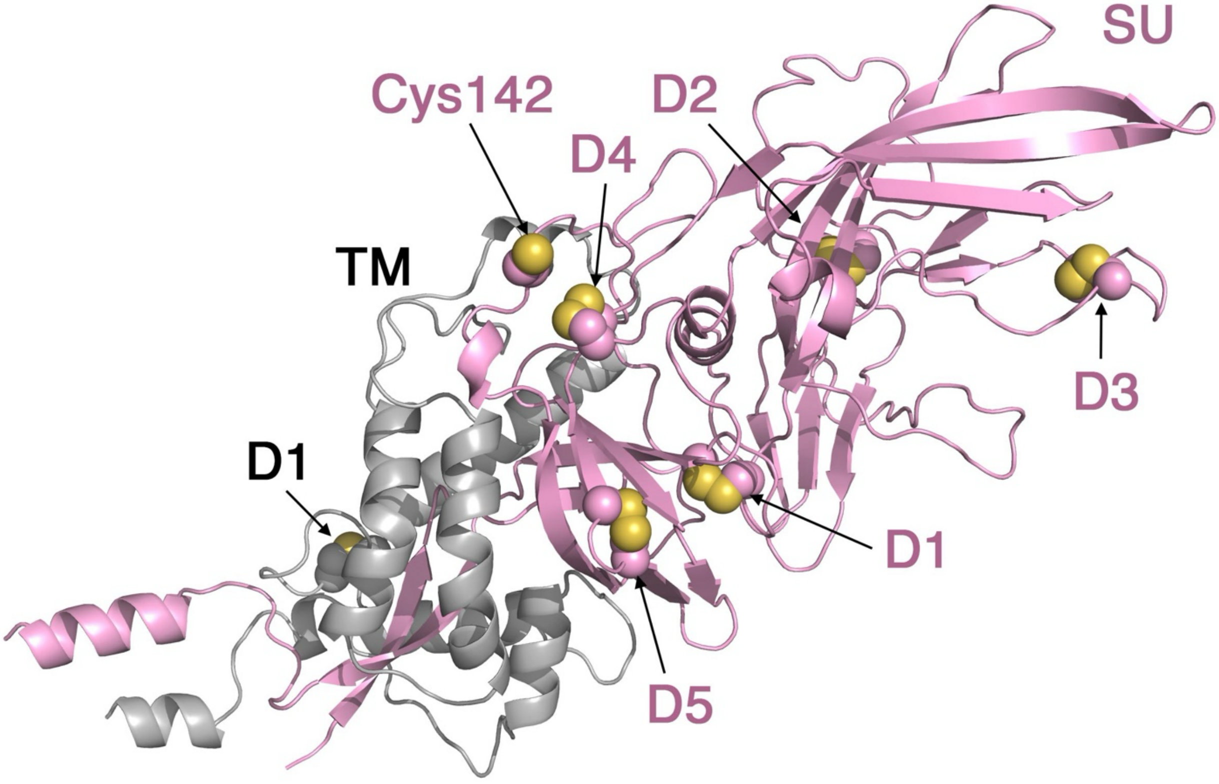


Figure S10 – **The disulfide bonds in the SU and TM domains**. A single SU (pink) and TM (grey) pair is shown as a ribbon diagram. The disulfide bonds are highlighted as spheres and noted, as well as the single unpaired cysteine residue in SU.

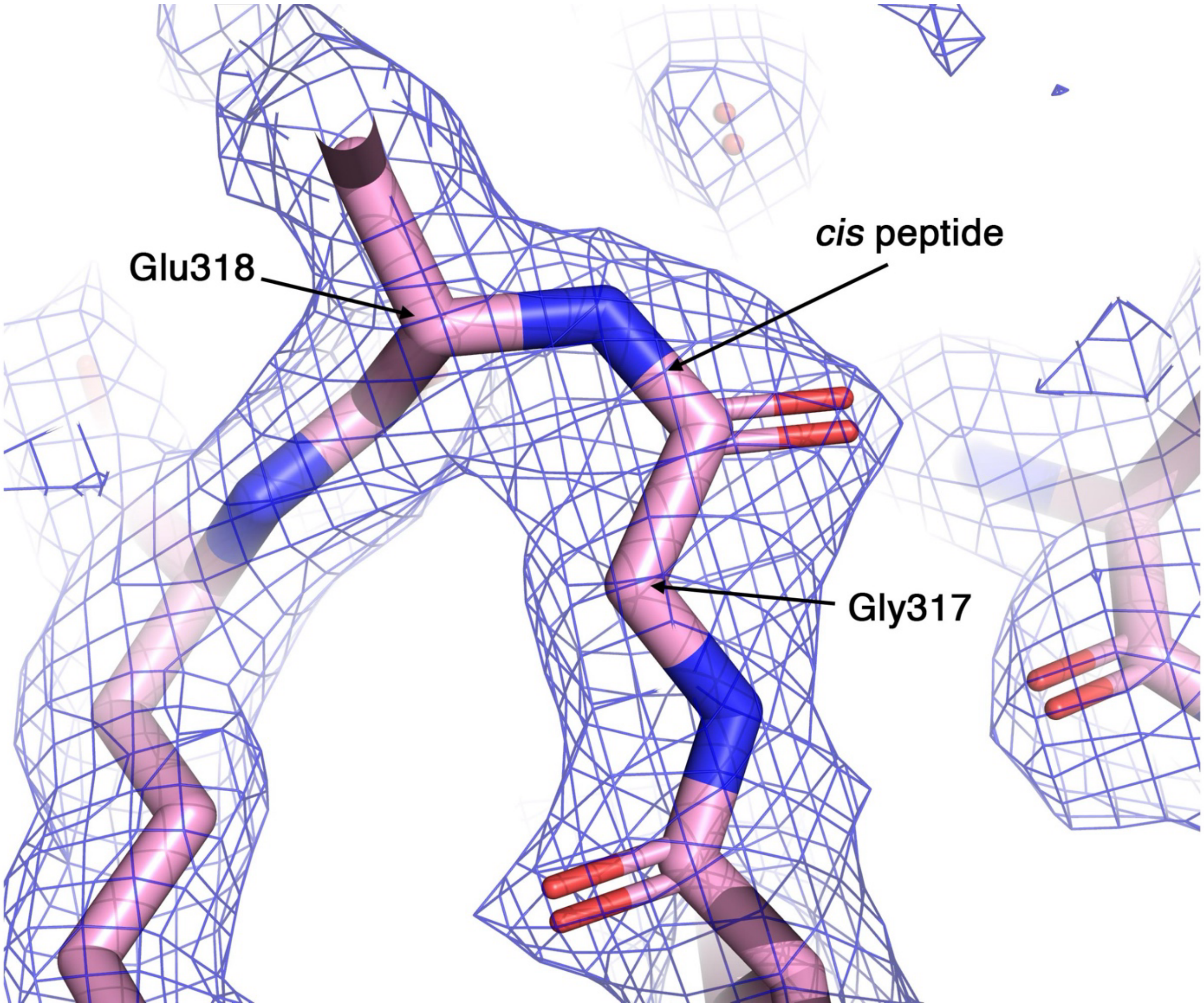


Figure S11 – **A cis-peptide in the SU domain**. A *cis* peptide configuration between Gly317 and Glu318 is shown. EM density at $\sigma$=5 is shown as blue mesh.

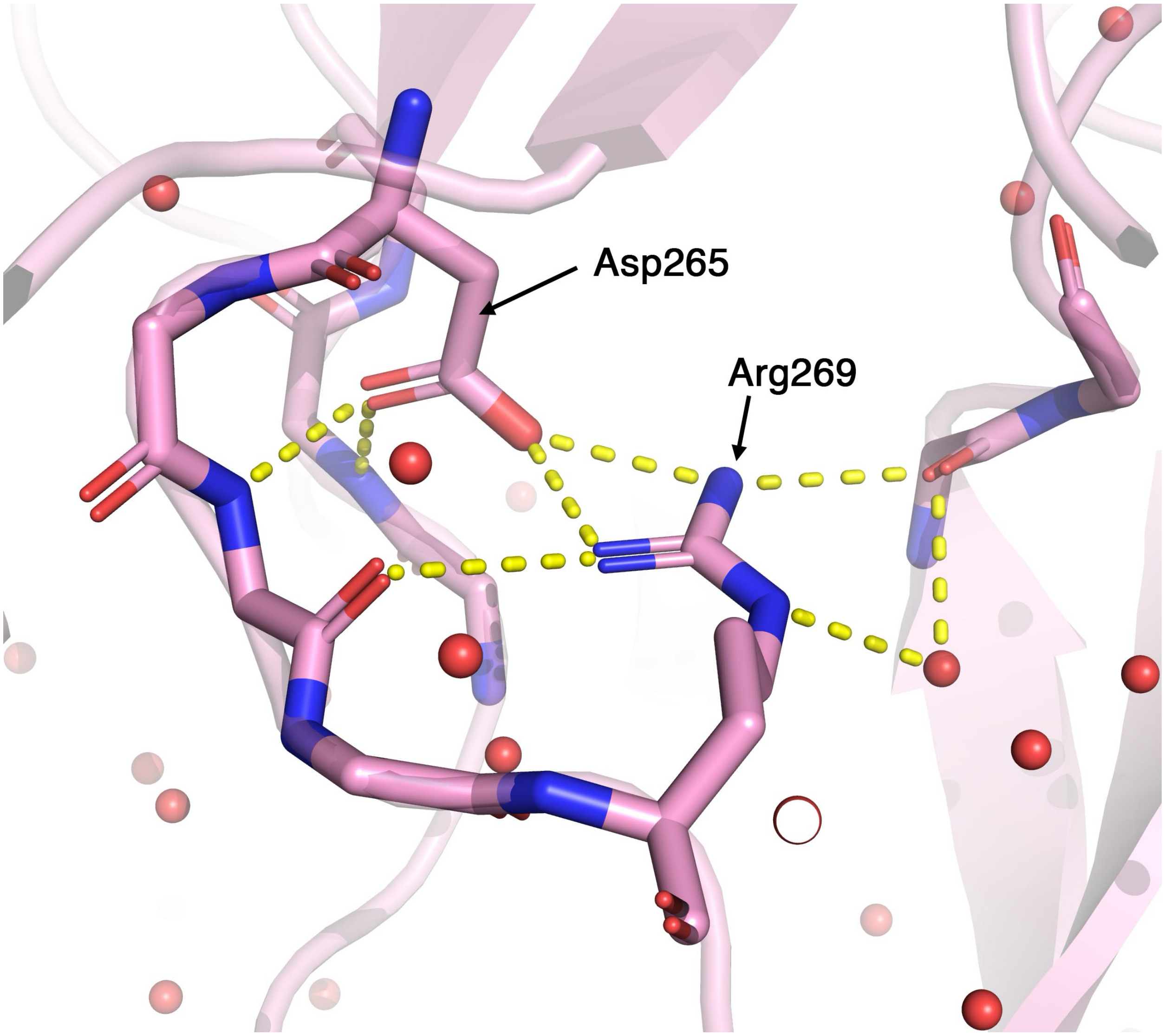


Figure S12 – **A salt bridge in the core of the SU domain**. A salt bridge between Arg269 and Asp 265 is noted. Polar interactions are illustrated with yellow dashed lines. Water molecules are shown as red spheres.

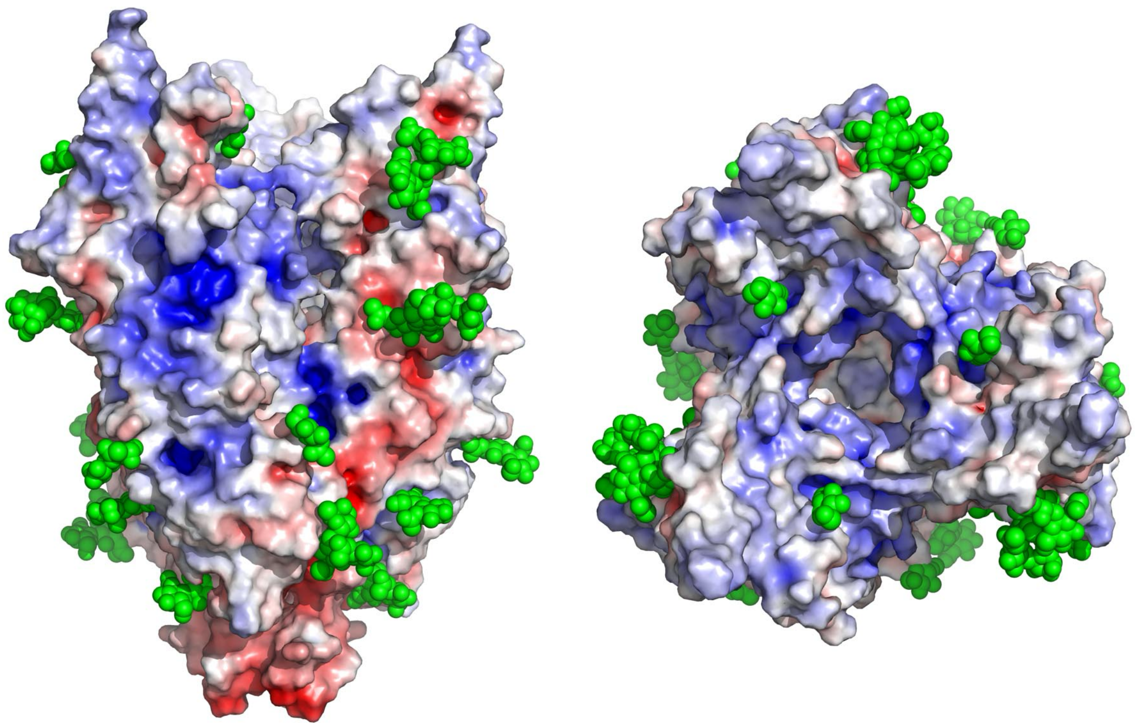

Figure S13 – **N-linked glycans and the putative binding sites for HS**. The HERV-K spike is shown with a surface electrostatic potential coloring, similar to the main Figure 4a. N-linked glycans are shown as green spheres. The spike is shown from a 'side' view and a 'top' view (left and right, respectively). Since only the proximal few sugars are visible for the glycan trees, their true size is not fully represented in this figure and is generally much bigger.

### Table S1 – Model refinement statistics

| Model | PDB 9NND | |
|---|---|---|
| **Composition (#)** | | |
| Chains | 12 | |
| Atoms | 14763 (Hydrogens: 0) | |
| Residues | Protein: 1542 | |
| Water | 1626 | |
| Ligands | BMA: 3 | |
| | NAG: 48 | |
| | FUC: 3 | |
| | MAN: 6 | |
| **Bonds (RMSD)** | | |
| Length (Å) (# > 4σ) | 0.003 (0) | |
| Angles (°) (# > 4σ) | 0.609 (0) | |
| MolProbity score | 1.22 | |
| Clash score | 4.20 | |
| **Ramachandran plot (%)** | | |
| Outliers | 0.00 | |
| Allowed | 2.09 | |
| Favored | 97.91 | |
| **Rama-Z (Ramachandran plot Z-score, RMSD)** | | |
| whole (N = 1530) | 0.49 (0.21) | |
| helix (N = 303) | 2.06 (0.30) | |
| sheet (N = 390) | 0.06 (0.26) | |
| loop (N = 837) | -0.12 (0.20) | |
| Rotamer outliers (%) | 0.00 | |
| Cβ outliers (%) | NA | |
| **Peptide plane (%)** | | |
| Cis proline/general | 7.9/0.2 | |
| Twisted proline/general | 0.0/0.0 | |
| CaBLAM outliers (%) | 0.40 | |
| **ADP (B-factors)** | | |
| Iso/Aniso (#) | 14763/0 | |
| min/max/mean | | |
| Protein | 0.29/99.59/22.86 | |
| Ligand | 10.46/84.45/40.61 | |
| Water | 1.00/55.29/24.60 | |
| **Occupancy** | | |
| Mean | 1.00 | |
| occ = 1 (%) | 100.00 | |
| 0 < occ < 1 (%) | 0.00 | |
| occ > 1 (%) | 0.00 | |
| **Data** | | |
| **Box** | | |
| Lengths (Å) | 98.88, 101.35, 131.84 | |
| Angles (°) | 90.00, 90.00, 90.00 | |
| Supplied Resolution (Å) | 2.1 | |
| Resolution Estimates (Å) | Masked | Unmasked |
| d 99 (full) | 2.2 | 2.2 |
| d model | 2.2 | 2.2 |
| d FSC model (0/0.143/0.5) | 1.8/2.0/2.2 | 1.8/2.1/2.2 |
| Map min/max/mean | -1.60/2.75/0.02 | |
| **Model vs. Data** | | |
| CC (mask) | 0.87 | |
| CC (box) | 0.78 | |
| CC (peaks) | 0.81 | |
| CC (volume) | 0.85 | |
| Mean CC for ligands | 0.68 | |
